# Functional connectivity alterations in Parkinson’s disease during the stop-signal task

**DOI:** 10.1101/304584

**Authors:** Chris Vriend, Douglas T. Leffa, James P. Trujillo, Niels J.H.M. Gerrits, Femke E. de Boer, Henk W. Berendse, Ysbrand D. van der Werf, Odile A. van den Heuvel

## Abstract

Although deficits in response inhibition are common in Parkinson’s disease (PD), studies on their neural correlates are relatively scarce. In our previous study, medication-naïve PD patients showed, among others, lower activation of the inhibition-related bilateral inferior frontal gyrus (IFG) compared with healthy controls while performing the stop-signal task. Here we report on a follow-up study in the same cohort.

Fourteen medicated PD patients and 16 healthy controls performed the same stop-signal task during MRI; 3.1±1.0 years after our previous study. During that time, all PD patients had started to use dopaminergic medication. We analyzed task-related functional activity and connectivity in the bilateral IFG and anterior insula, brain regions that are important response inhibition. Task-related functional connectivity was analyzed with generalized psychophysiological interaction.

PD patients were significantly slower on response initiation (GO reaction time) and response inhibition (stop-signal reaction time) than healthy controls. There were no between-group differences in functional activity. On the other hand, functional connectivity of the IFG and anterior insula was significantly lower in PD compared with healthy controls. Mainly the inferior parietal lobule and dorsolateral prefrontal cortex were less connected with these seed regions.

These results show that early-stage medicated PD patients show lower task-related functional connectivity but not activity of brain regions that are important for response inhibition; the IFG and anterior insula. We tentatively speculate that the use of dopaminergic medication upheld task-related activity but not connectivity.

## 1. Introduction

Response inhibition is an experimental operationalization of impulse control [1] and can be used to study impulsivity in Parkinson’s disease (PD). Response inhibition tasks may also aid in monitoring and predicting disease progression, including cognitive decline [2]. Compared with healthy subjects, PD patients generally show behavioral impairments on tasks that measure response inhibition, particularly during more demanding tasks, such as the stop-signal task [3, 5].

Neuroimaging studies on response inhibition in PD patients are still relatively scarce. Studies in medicated PD patients have shown higher activation of prefrontal and striatal brain areas during a Go/No-Go task compared with healthy controls [6, 7] and lower activation of the inferior frontal gyrus (IFG) and pre-supplementary motor cortex during the stop-signal task [4, 8]. Recently, we studied the stop-signal task during functional Magnetic Resonance Imaging (fMRI) in early-stage, untreated PD patients to avoid the confounding influence of dopaminergic medication [9]. We showed that these de novo PD patients, when compared with healthy controls, showed normal behavior but, among other, lower task-related activation of the bilateral IFG.

Here we report on a follow-up study (3.1±1.0 years) in the abovementioned cohort of previously medication-naïve PD patients and control subjects, who performed the stop-signal task during MRI scanning. Because of a scanner upgrade in between the study at baseline [9] and the current study, we were unable to perform a longitudinal analysis of the imaging data. We therefore performed a cross-sectional analysis of the imaging data at the follow-up measurement only. We chose the bilateral IFG and anterior insula as regions of interest (ROI), because we previously reported lower task-related activation of the IFG in PD patients [9] and the anterior insula has recently been shown to be vital for response inhibition [10] and highly vulnerable to accumulation of alpha-synuclein in PD [11]. Based on the results of our previous study in this sample, we hypothesized that PD patients would show lower task-related activation of these brain areas. Brain areas do not work in isolation but work together to instigate behavior. We therefore additionally performed task-related functional connectivity analyses, based on generalized psychophysiological interaction (gPPI), with the left and right IFG and anterior insula as seed regions. GPPI allows for the analysis of functional connectivity between brain regions in the context of a specific task by adding psychophysiological interactions as regressors to the model [12]. Based on our own previous gPPI studies in this sample [13] and studies from others [14, 15], we further hypothesized that task-related functional connectivity during the stop-signal task would be lower in PD patients compared with well-matched healthy subjects.

## 2. Methods

### 2.1 Participants

Participants were recruited as part of a follow-up study in our previously established group of 21 non-demented medication-naïve PD patients and 37 matched healthy controls. Sixteen PD patients and 16 healthy controls accepted our invitation for re-assessment after a mean follow-up period of 3.1 ± 1.0 years. Subjects that were included in the current study and those that were lost to follow-up did not differ on demographic, clinical or behavioral data at baseline (data not shown). At follow-up, all PD patients still met the UK Parkinson’s disease Brain Bank criteria for idiopathic PD [16] and had started to use dopamine replacement therapy. Total levodopa equivalent daily dosage (LEDD) scores were calculated [17]. We used the Unified Parkinson’s Disease Rating Scale motor section (UPDRS-III) and modified Hoehn and Yahr stage to assess disease severity and disease stage, respectively. We used the Questionnaire for Impulsive-Compulsive disorders in Parkinson’s disease Rating Scale (QUIP-RS) to rate the severity of symptoms of impulse control disorders (ICD) (including QUIP-RS≥10 as cut-off for clinical relevant signs of ICD). According to the Mini-Mental State Examination (MMSE), none of the participants showed signs of dementia (all MMSE-scores > 24). None of the participants met our exclusion criteria based on excessive movement during MRI scanning (>3mm) or the use of psychotrophic drugs. All participants provided written informed consent according to the declaration of Helsinki, and the study was approved by the local research ethics committee (2008/145). All measurements were conducted while PD patients were in their ON state.

### 2.2 Stop-signal task

Details on the visual stop-signal task during fMRI scanning are described in our previous publication [9]. Briefly, while in the scanner, participants had to respond to in what direction (left or right) an arrow on the screen was pointing (Go-trials) using both hands. Twenty percent of all trials were stop-trials, signaled by a delayed presentation of a cross superimposed on the arrow. During those trials, participants had to refrain from responding. The delay of the stop-signal (stop-signal delay or SSD) was continuously adapted by a staircase tracking mechanism to approximate a 50% successful inhibition on all stop-trials. We measured the mean reaction time on successful Go-trials, the mean stop-signal reaction time (SSRT) and the error percentage on Go-trials and Stop-trials. Participants with a Go-trial error percentage >40% were excluded [18]. We used the integration method to estimate the SSRT [19]. Because response latencies gradually increased during the task, SSRTs were estimated separately in four smaller blocks (each block consisting of at least 50 trials) and subsequently averaged. Blocks with stop-trial error percentages <25% or >75% were excluded [18].

### 2.3 Image acquisition

MRI scans were acquired on a Discovery* MR750 (General Electric) scanner, at VU university medical center (Amsterdam, The Netherlands) in 2014. The participants’ heads were immobilized using foam pads to reduce motion artifacts. T2*-weighted echo-planar images (EPI’s) with blood oxygenation level-dependent (BOLD) contrast were acquired in each session; TR= 2100 ms, TE = 29 ms, flip angle=80°, 40 slices (3.75 × 3.75 mm in-plane resolution; 2.8 mm slice thickness; matrix size 64 × 64) per EPI volume. Structural images were acquired using a 3D sagittal T_1_-weighted sequence (TI=450 ms, TE=3 ms, voxel size 1 × 0.977 × 0.977 mm, 172 slices).

### 2.4 Behavioral and clinical data analysis

All between-group differences in demographic and clinical variables, except IQ estimation, were analyzed with two-sample T-tests or Mann-Whitney U-tests, depending on the distribution. We analyzed time effects on MMSE and UPDRS-III score with the Wilcoxon Signed Rank test. Analyses were conducted in IBM SPSS 22 (Armonk, NY, USA). Behavioral performance (i.e., SSRT and GO-RT) was analyzed with two-sample T-tests. Spearman’s rho correlations (r_s_) were performed between the behavioral and clinical measures. The statistical threshold for analyses on the behavioral and clinical measures was set to P<.05.

### 2.5 Data preprocessing and functional activity analyses

Imaging preprocessing was performed with SPM8 (Wellcome Trust Center for Neuroimaging, London, UK). Functional brain images were slice-time corrected, co-registered to the structural T1 image, normalized to Montreal Neurological Institute (MNI) space and spatially smoothed using an 8 mm Full-Width-at-Half-Maximum (FWHM) Gaussian kernel. All scans were visually inspected for possible motion artifacts. Successful Go-trials, successful Stop-trials and unsuccessful Stop-trials were modeled per participant using delta functions convolved with the hemodynamic response function. We additionally applied a 128 s high-pass filter and added six movement parameters as regressors of no interest. Inhibition-related BOLD activity was modeled by contrasting successful Stop-trials with successful Go-trials (Successful Stop>Successful Go) and brought to second-level random effects analyses.

This study aimed to follow up on the inhibition-related hypoactivation of the left and right IFG that we found in the de novo state of these PD patients [9]. We therefore examined group effects in the same 10-mm spherical ROIs in the bilateral IFG as previously reported (right IFG: x=51, y=20, z=7; left IFG: x=-54, y=17, z=4). Based on a recent meta-analysis we additionally selected the left (x=-40, y=16, z=-4) and right (x=34, y=22, z=-4) anterior insula as ROIs, because the anterior insula is one of the most consistently activated brain regions during the stop-signal task [10]. The coordinates of the anterior insula were derived from this meta-analysis. We used MarsBaR (http://marsbar.sourceforge.net) to construct all four ROIs. Voxels that showed overlap between the insular and IFG ROIs were excluded from the final ROIs. Imaging data were analyzed using two-sample T-tests and masked by the main effects of the task to ensure that results are specific to differences in task activation. The statistical threshold was set to P<.05, FWE-corrected for the ROI analyses. We also report exploratory whole-brain analyses at a more lenient threshold of P<.001, uncorrected, k_e_ ≥ 5, in the supplements.

### 2.6 Functional connectivity

We additionally examined task-related functional connectivity between the four seed regions - the bilateral IFG and anterior insula - and the rest of the brain in a voxel-wise manner. We used McLaren’s generalized form of context-dependent psychophysiological interaction (gPPI; http://www.nitrc.org/projects/gppi) to model all psychological task conditions into one first-level design [12]. We used the same 10 mm spheres as seed regions as we did for the BOLD activity ROI analyses. In the gPPI analysis, time-series extraction from the seed region was restricted to voxels within the subject-specific activation mask. First-level models consisted of 13 regressors: three convolved task regressors (successful Go-trials, successful Stop-trials and unsuccessful Stop-trials), their three convolved PPI regressors, one regressor for the time-series of the seed-region and six movement parameters. We contrasted the successful Stop-trials PPI with successful Go-trials PPI (Successful Stop PPI >Successful Go PPI). At second level, between-group effects on functional connectivity were analyzed separately for each seed-region with two-sample T-tests. Statistical threshold for all four whole-brain analyses (one per seed) was set to P<.001, uncorrected with a cluster size threshold of k_e_≥5. All contrasts were masked by the group-specific main effects of task (inclusive mask, *P*<.05).

## 3. Results

### 3.1 Demographic and clinical analyses

Two PD patients had to be excluded from analyses; one due to technical difficulties during scanning, the other because of a too high error percentage on the GO-trials (>40%), leaving 14 PD patients and 16 healthy controls for analyses. Table 1 summarizes the sample characteristics. Groups were matched on age, gender and education.

**Table 1.**
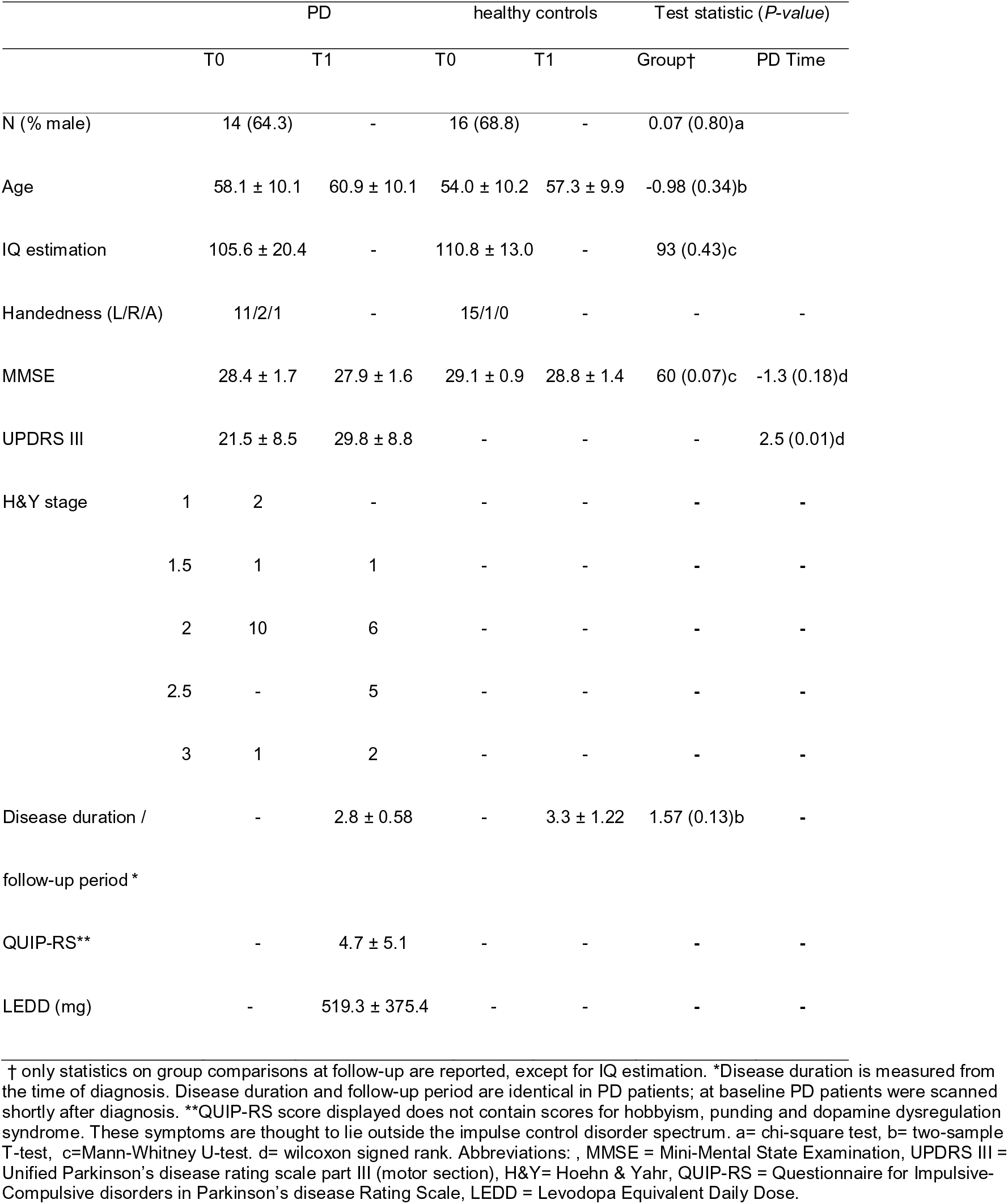
– Sample characteristics, clinical and behavioral data

### 3.2 Behavioral analysis

Behavioral analysis showed that both GO-RT (*t*(_28_)=-2.2, *P*=.04) and SSRT were significantly slower in PD patients compared with healthy controls (*t*(_28_)=-2.40, P=.02). Behavioral performance did not correlate with motor symptom severity or severity of ICD symptoms.

### 3.3 BOLD activity analyses

Analyses on task-related BOLD response BOLD activity analysis revealed no between-group differences in inhibition-related activity, neither in the ROI analysis nor the exploratory whole-brain analysis (see supplements).

### 3.4 Functional connectivity analyses - between group differences

Compared with healthy controls, PD patients showed lower functional connectivity between all seed regions (bilateral IFG and anterior insula) and the IPL. Furthermore, the left anterior insula showed lower connectivity with, amongst others, the dorsolateral prefrontal cortex (DLPFC), while the right IFG showed lower connectivity with the precuneus and DLPFC. See Figure 1 and Table 2 for a complete overview of the between-group differences.

**Figure 1.**
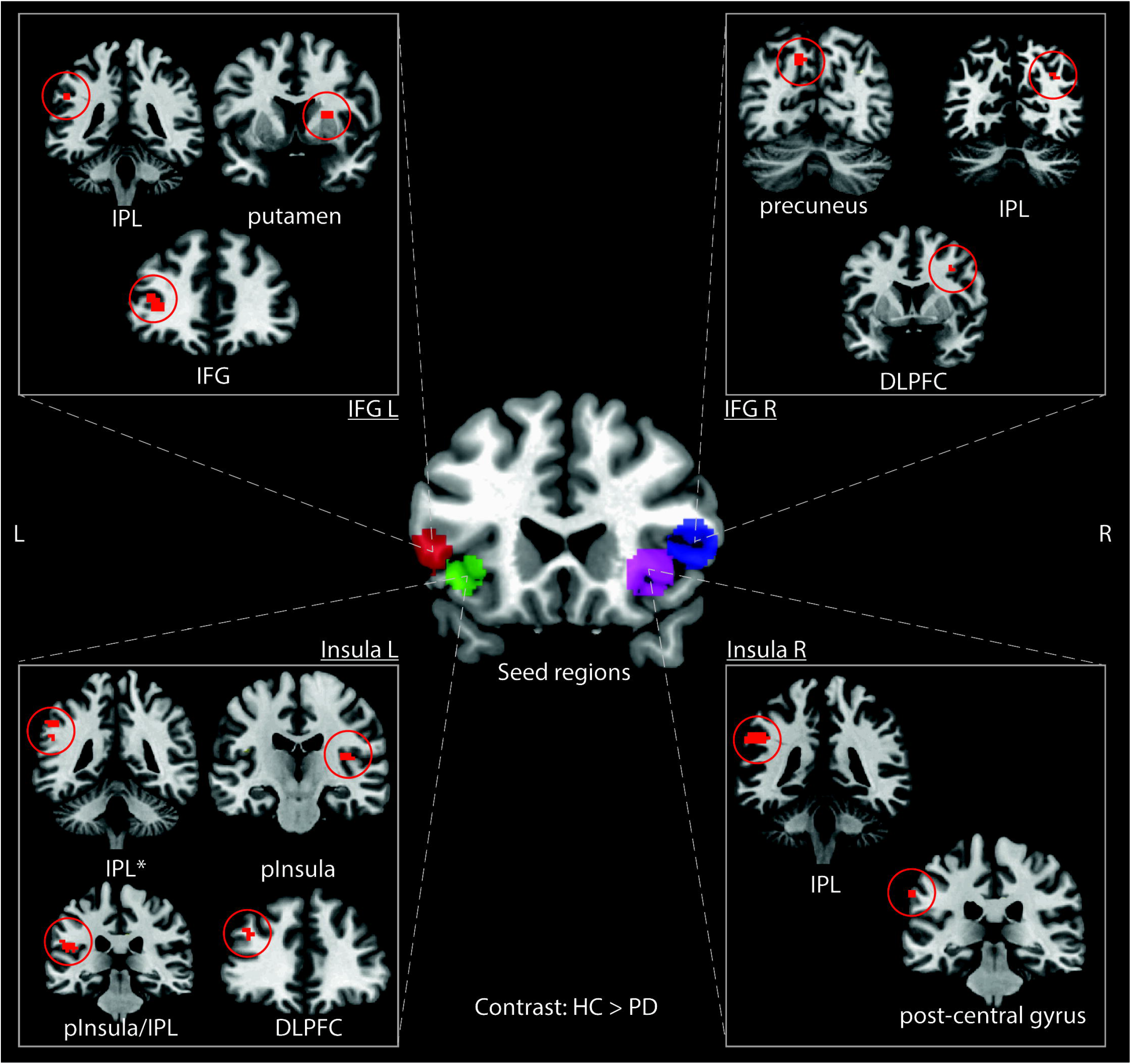
Functional connectivity analysis. The coronal brain image in the middle of the figure shows the placement of the seed regions in the left anterior insula (green), left inferior frontal gyrus (red), right anterior insula (purple) and right inferior frontal gyrus (blue). Between-group differences (HC>PD) in coupling between these seed regions and clusters in other brain regions are depicted in the white boxes. Left in the brain is left on the coronal brain image. See Table 2 for the MNI coordinates of these significant clusters. Abbreviations: PD = Parkinson’s disease, HC = healthy control, IPL = inferior parietal lobule, IFG = inferior frontal gyrus, DLPFC = dorsolateral prefrontal cortex, pInsula = posterior insula. *= only one of clusters in the IPL that showed significantly lower coupling in PD is shown (see Table 2).

**Table 2.** - functional connectivity – between-group effects

| *Contrast: HC>PD* |  |  |  |  |  | MNI coordinates |  |  |
| --- | --- | --- | --- | --- | --- | --- | --- | --- |
| Seed | Anatomical region | BA | L/R | $k_e$ | $t$ | x | y | z |
| *Insula R* | Postcentral gyrus | 2 | L | 11 | 4.71 | -66 | -28 | 37 |
|  | IPL | 40 | L | 36 | 4.51 | -48 | -40 | 34 |
| *Insula L* | Posterior insula/IPL | 13/40 | L | 39 | 4.50 | -51 | -28 | 19 |
|  | Posterior insula | 13 | R | 14 | 4.03 | 36 | -19 | 16 |
|  | IPL | 40 | L | 34 | 4.26 | -54 | -43 | 40 |
|  |  |  | L | 9 | 4.23 | -42 | -52 | 37 |
|  |  |  | L | 5 | 4.06 | -66 | -40 | 22 |
|  | DLPFC | 46 | L | 5 | 3.76 | -39 | 38 | 28 |
| *IFG R* | Precuneus | 7 | L | 12 | 4.17 | -12 | -67 | 43 |
|  | Precuneus/IPL | 7/40 | R | 6 | 3.82 | 33 | -67 | 34 |
|  | DLPFC | 9 | R | 5 | 3.81 | 36 | 5 | 37 |
| *IFG L* | IFG | 47 | L | 28 | 5.33 | -36 | 41 | 4 |
|  | Putamen |  | R | 20 | 4.69 | 24 | 8 | 13 |
|  | IPL | 40 | L | 5 | 4.33 | -51 | -40 | 34 |

$P < .001$ , uncorrected, $k_e \geq 5$ . To preserve space, contrast $PD > HC$ , without any significant findings, has been omitted from the table. Abbreviations: PD = Parkinson's disease, HC=healthy controls, BA=Brodmann Area, $K_e$ = cluster size, IPL= inferior parietal lobule, DLPFC= dorsolateral prefrontal cortex, IFG=inferior frontal gyrus.

### 4. Discussion

This study is a follow-up on our previous fMRI study with the stop-signal task where PD patients were recently diagnosed and still naïve for dopaminergic medication [9]. In our previous study we observed comparable behavioral performance but lower task-related activation of the bilateral IFG and IPL in PD patients compared with healthy controls. Consistent with other studies, the current study showed that PD patients perform worse on the stop-signal task compared with healthy controls [3, 4, 6, 7]. Between-group differences in task-related activation were, however, not observed. All PD patients had started to use dopaminergic medication after participation in the previous study and were all on medication during scanning. We tentatively postulate, consistent with the ON/OFF Go/No-Go study from Farid et al. [7], that the use of dopaminergic medication at follow-up normalized task-related activity in the PD patients compared with our healthy controls by (partially) restoring dopamine concentrations to physiological levels [20]. Nevertheless, Ye et al. 2014 showed that even on medication, PD patients show reduced activation of the IFG compared with healthy controls. Although the sample from Ye et al. and ours are quite comparable with respect to motor symptom severity and disease stage, patients in our sample had a much shorter disease duration (2.8 versus 10.8 years), were slightly younger and had a lower LEDD. As dopaminergic medication loses its potency with progression of the disease [21] and loses some of its beneficial effect on behavioral performance of response inhibition tasks [22], it is possible that dopaminergic medication normalized task-related activity in our sample but not that of Ye et al. 2014.

In contrast to our functional activation analyses, our seed-based functional connectivity analyses - using gPPI - revealed multiple between-group differences. In PD patients, all four seeds showed lower connectivity with the IPL, compared with healthy controls. The IPL is consistently activated in tasks involving cognitive control [10] and part of a cognitive control network that subserves executive functions, including response inhibition [23]. Although we previously showed a lower activation of the IPL during the task while PD patients were still de novo [9], task-related activation of the IPL did not differ in the follow-up sample (data not shown), and neither did task-related functional connectivity of the bilateral IFG and anterior insula in the sample at baseline (data not shown). PD patients additionally showed lower connectivity between the seed regions and the DLPFC and precuneus. The IPL, DLPFC and precuneus are all associated with the cognitive control network, suggesting a general lower connectivity of the cognitive control network in PD during response inhibition.

Our seed regions are situated in the fronto-insular region, a region that is critically involved in switching activation of brain networks [24] and prone to accumulation of alpha-synuclein when PD progresses from Braak stage III to V [11]. Seed-based analyses of task-related functional connectivity are therefore likely to show desynchronization of these seed regions with other brain areas with progression of PD. Our results are also consistent with our previous finding of lower connectivity between the DLPFC and IPL in the same sample of medicated PD patients during planning in the Tower of London Task [13].

Working under the tentative assumption that the use of chronic dopaminergic medication by our PD patients at follow-up led to comparable task-related *activation* with healthy controls, dopaminergic medication did not seem to have the same influence on functional *connectivity*. There is some evidence from ON/OFF paradigms suggesting that levodopa reduces, instead of enhances, functional connectivity relative to both the OFF phase and matched healthy controls [25, 26]. Dopaminergic medication might thus have a dissociable effect on functional activity and connectivity. Dopaminergic medication also has differential effects on the functional connectivity of the various resting-state networks [27] and depends on PD motor subtype [28]. Because we were unable to perform longitudinal within-subject analyses and did not assess our patients ON and OFF medication, the between-group differences on task-related functional activation versus functional connectivity warrants further investigation.

A number of limitations have to be borne in mind while interpreting these results. Unfortunately, during the follow-up period the MRI scanner used for the baseline measurements had to be serviced and upgraded, limiting our ability to simultaneously model group and time effects in a full factorial analysis to study the task-related activity and connectivity longitudinally. Another limitation stems from the fact that a number of our patients was lost to follow up due to exacerbation of PD progression or other comorbidities, possibly leading to a selection bias. Nevertheless, demographic, clinical and behavioral data at baseline of subjects included in the follow-up analysis and those that were lost to follow-up were comparable.

In conclusion, using a stop-signal task during fMRI imaging we showed lower task-related functional connectivity among brain areas within the cognitive control network in medicated PD patients compared with matched healthy controls, but no differences in activation of these brain areas; i.e. the bilateral IFG and anterior insula. This contrasts with our previous study in this sample that showed hypoactivation of the bilateral IFG when patients were recently diagnosed and still medication-naïve [9]. We tentatively presume a role of dopaminergic medication at follow-up in upholding task-related activity but not connectivity but this warrants further investigation using on/off paradigms.

## Acknowledgement

This study was supported by a grant from the Dutch Parkinson Patient Association and Stichting Parkinson Fonds.

